# Serotonergic signaling governs *C. elegans* sensory response to conflicting olfactory stimuli

**DOI:** 10.1101/2025.03.19.644218

**Authors:** Caroline S. Muirhead, Sophia Guerra, Bennett W. Fox, Frank C. Schroeder, Jagan Srinivasan

## Abstract

Neural circuits that consolidate sensory cues are essential for neurological functioning. Neural circuits that perform sensory integration can vary greatly because the sensory processing regions of the brain employ various neural motifs. Here, we investigate a neural circuit that mediates the response to conflicting olfactory stimuli in *C. elegans*. We concurrently expose animals to an aversive dispersal pheromone, osas#9, and an attractive bacterial extract. While worms usually avoid osas#9 alone, they suppress this avoidance behavior in the presence of a bacterial extract. Loss-of-function mutants and cell-specific rescues reveal that serotonergic signaling from the ADF neuron is essential for bacterial extract-induced osas#9 avoidance attenuation. The inhibitory serotonin receptor, MOD-1, which is widely expressed on interneurons and motor neurons, is required for this sensory integration, suggesting that serotonin acts in an inhibitory manner. By performing calcium imaging on the ADF neurons in synaptic signaling (*unc-13*) and peptidergic (*unc-31*) signaling mutant backgrounds, we show that the ADF neurons require input from other neurons, likely the ASK neurons, to respond to food extracts. We reveal a cue integration neural circuit in which serotonergic signaling at the sensory neuron level silences an aversive neural signal.

**Significance:** Animals use sensory cues to make behavioral choices and sometimes, these cues convey opposite information. The nervous system consolidates competing sensory cues to create a coherent response to external stimuli. The neural circuits that govern this process are important, and still largely unknown. We use *C. elegans,* a soil-dwelling nematode, to uncover a neural circuit governing the consolidation of competing cues by concurrently exposing worms to positive and negative stimuli . We find that the neurotransmitter serotonin can suppress aversive neural signals created by negative stimuli. These results show the important neurological role that serotonin plays in modulating neural signals.

## Introduction

The nervous system’s ability to consolidate sensory input is essential (Wallace et al., 2020). The nervous system constantly receives sensory input from multiple sensory modalities (Stein et al., 2014; Yau et al., 2015). Additionally, the nervous system must consolidate competing sensory information from the same modality (Blake and Logothetis, 2002; Cohen Kadosh et al., 2008). Impairment in sensory integration is associated with a variety of neurological disorders, including autism spectrum disorder, attention deficit hyperactivity disorder, and schizophrenia (Ghanizadeh, 2011; Koziol et al., 2011; Tseng et al., 2015). Across the animal kingdom, neural circuits that perform sensory integration tasks generally use coordinated excitation and inhibition (Yan, 2002; Wang, 2008; Miller et al., 2011; Wu and Guo, 2011; Chadderton et al., 2014; Herd et al., 2021). In the human brain sensory information processing is complex, and therefore, these circuits can vary (Bertrand and Thomas, 2004; Chadderton et al., 2014; Leighton and Lohmann, 2016). In fact, models of sensory integration in mammals are only made to apply to local neural circuits – not the whole mammal brain (Sabes, 2011).

*C. elegans* are an ideal model for studying small neural circuits that integrate sensory information. They are both simple and amenable to experimentation (Chalfie and Jorgensen, 1998). *C. elegans* hermaphrodites contain precisely 302 fully mapped neurons (White et al., 1986). Additionally, *C. elegans* display robust behavioral outputs in response to various stimuli (de Bono and Maricq, 2005) and since they are transparent, observing fluorescent protein expression and imaging experiments are straightforward (Félix and Braendle, 2010). Together, these attributes allow for manipulation of specific neurons, receptors, or neural connections and observation of the corresponding behaviors, making it possible to understand how individual cells act within a neural circuit.

Since *C. elegans* lack complex sensory organs, they communicate via olfaction (Bargmann, 2006). Pheromones released by *C. elegans*, ascarosides, are used to communicate a wide variety of information (Ludewig and Schroeder, 2013; Muirhead and Srinivasan, 2020); worms will attract mates (Izrayelit et al., 2012; Chute and Srinivasan, 2014), communicate food levels (Artyukhin et al., 2013), and promote entry into dauer phase using these secreted pheromones (Albert et al., 1981; Edison, 2009). Here, we utilize a specific ascaroside, osas#9, to study sensory integration. osas#9 is produced primarily by food-deprived L1 animals (Artyukhin et al., 2013). Given that the biosynthesis of osas#9 is dependent on starvation, it is unsurprising that it functions as a dispersal cue and communicates lack of food in the environment (Chute et al., 2019). Structurally, osas#9 is unique among the ascarosides because it incorporates the neurotransmitter octopamine (Artyukhin et al., 2013; Chute et al., 2019). Previous work with osas#9 has determined that it is sensed primarily by the nociceptive ASH sensory neuron via a GPCR, TYRA-2 (Chute et al., 2019).

Here, we concurrently exposed animals to osas#9 (an aversive cue) with an opposite cue (an attractive bacterial extract) to determine how the nervous system reconciles competing olfactory cues. We find that serotonergic signaling from the ADF sensory neurons is essential for the integration of the food cue into the nervous system. We also show that the ADF neurons act with other neurons in order to achieve this integration.

## Methods

### Worm maintenance

Animals were maintained on nematode growth media (NGM) plates at 20°C. Animals were fed OP50 *E. coli* (Brenner, 1974); young adult animals were transferred to new plates to avoid starvation. Worm strains used are shown in table S1.

### Avoidance assay

The avoidance assay was performed as previously described (Hilliard et al., 2002). Animals were transferred onto unseeded NGM plates. Animals were starved one hour prior to behavioral testing. A copper ring was placed on the NGM plate to minimize animals leaving the plate.

The avoidance assay was performed by placing a small (roughly 5nL) drop of stimulus dissolved in diH2O on the tail of a forward moving animal. Drops were created using capillary tubing. Once the liquid encountered the animal’s head, the animal’s response would be scored as avoidance or non-avoidance. If within five seconds of encountering a drop, the worm showed at least two body bend reversals or a 180° omega turn, the behavior was scored as avoidance (Gray et al., 2005). Other responses (continued forward movement, stopping without a reversal, or one backward body bend) were scored as non-avoidance. Animals were scored blindly with respect to genotype. 8-15 individual worms were tested per plate and a minimum of six plates were tested over a minimum of three different days. For each individual plate, an avoidance index was calculated. Avoidance index was calculated as number of worms that avoid a stimulus divided by the total number of worms tested per plate.

### Calcium imaging

Prior to imaging, worms were starved for 1hr on NGM plates so that they would have the same internal state during imaging and behavior.

Calcium imaging was performed in a microfluidic device previously described by Reilly et al (Reilly et al., 2017). Animals were immobilized in a PDMS chip. The nose of the worm was exposed to either solvent control or stimulus liquid. All stimuli were dissolved in S. basal buffer with 1mM tetramisole as a paralytic to assist in immobilizing the animals. Solvent and flow controls were 1mM tetramisole S. basal with 0.1µg/mL and 0.2µg/mL of fluorescein, respectively.

During a trial, an animal was exposed to the solvent control for five seconds. Then, it was exposed to a pulse of stimulus for 30 seconds. Finally, it was exposed to solvent control for another 15 seconds.

Intensity of pixel fluorescence in neurons was analyzed using Fiji. ΔF was calculated by subtracting the fluorescence intensity of the background from the fluorescence intensity of the neuron body. Fo is the average ΔF for ten frames prior to stimulation. To obtain %Δ F/Fo, one was subtracted from ΔF/Fo, then that quantity was multiplied by 100. Max peak ΔF/Fo refers to the largest ΔF/Fo value detected.

### Microscopy

Animals were mounted on a 2% noble agar pad and immobilized using 1µL of 1M sodium azide. Images were obtained using a Zeiss LSM510 Meta inverted confocal microscope. 1µm thick z-stacks slices were used to capture the whole head region of the animal. Images were analyzed using ImageJ software.

### Preparation of bacterial extract

10 mL of methanol was added to one gram of freeze-dried *E. coli* OP50 powder (InVivoBiosystems, formerly NemaMetrix Inc., OP-50-31772) in a 50 mL conical tube and then sonicated for 5 min (2 s on/off pulse cycle at 90 A) using a Qsonica Q700 Ultrasonic Processor with a water bath cup horn adaptor (Qsonica 431C2). Following sonication, the conical tube was centrifuged (3000 x *g*, 22 °C, 5 min) and the resulting clarified supernatant transferred to a clean pear flask. This process was repeated twice more, and each time the clarified supernatant was transferred to the same pear flask. After the third extraction, the insoluble material was washed with 10 mL of methanol and vortexed for 10 s, then subject to centrifugation (3000 x *g*, 22 °C, 5 min) and the resulting clarified supernatant transferred to the same pear flask. One additional methanol wash was performed, and the supernatant was combined in the same pear flask, resulting in ∼ 50 mL methanol. An aliquot of the methanol extract was transferred to a clean 8 mL glass vial and concentrated to dryness in an SCP250EXP Speedvac Concentrator coupled to an RVT5105 Refrigerated Vapor Trap (Thermo Scientific). The resulting powder (14.6 mg) was resuspended in 1mL of methanol.

### Statistical analysis

Statistical analyses were performed in GraphPad Prism. When multiple comparisons were being made, the Shapiro-Wilk normalcy test was performed to determine if data were normally distributed. If data were normally distributed, an ordinary one-way ANOVA was performed followed by Sidak’s multiple comparisons. If data were not normally distributed, then a Kruskal-Wallis test was performed followed by Dunn’s multiple comparisons. When comparing an animal to itself (pre and post imaging stimulus) a Shapiro-Wilk normalcy test was performed. If the data were not normally distributed, a two-tailed Wilcoxon test was performed. Otherwise, a two-tailed paired t-test was performed.

## Results

### Food-deprived *C. elegans* suppress osas#9 avoidance in the presence of food odor

Previous work has demonstrated that food-deprived worms avoid the pheromone osas#9, whereas fed animals show no aversive behavior (Artyukhin et al., 2013; Chute et al., 2019)(Figure S1). Because of the myriad changes that occur between the fasted and fed states (Baugh and Hu, 2020), we wanted to create a situation in which worms must choose between two competing stimuli without providing a nutritive source. To do this, we concurrently exposed starved animals to both osas#9 and an extract of the bacterial food (*E.* coli) using the avoidance assay (Figure 1A) and observed that worms no longer avoided osas#9 in the presence of *E. coli* extract (Figure 1B). Since sensory processing is typically dependent on cue strength, we tested avoidance behavior to osas#9 mixed with varying food extract concentrations and saw that food extract-dependent osas#9 avoidance attenuation is concentration dependent (Figure 1C). We observed that high concentrations of food extract strongly suppressed avoidance to osas#9 (1/10- 1/2000) (Figure 1C). Conversely, worms avoided the osas#9 *E. coli* extract mixture at low concentrations, (1/5000-1/10000) (Figure 1C). We also observed that at intermediate concentrations of extract (1/3000-1/4000), worms did not avoid the mixture significantly more than the solvent control, but there was a trend of increased avoidance (Figure 1C).

**Figure 1:**
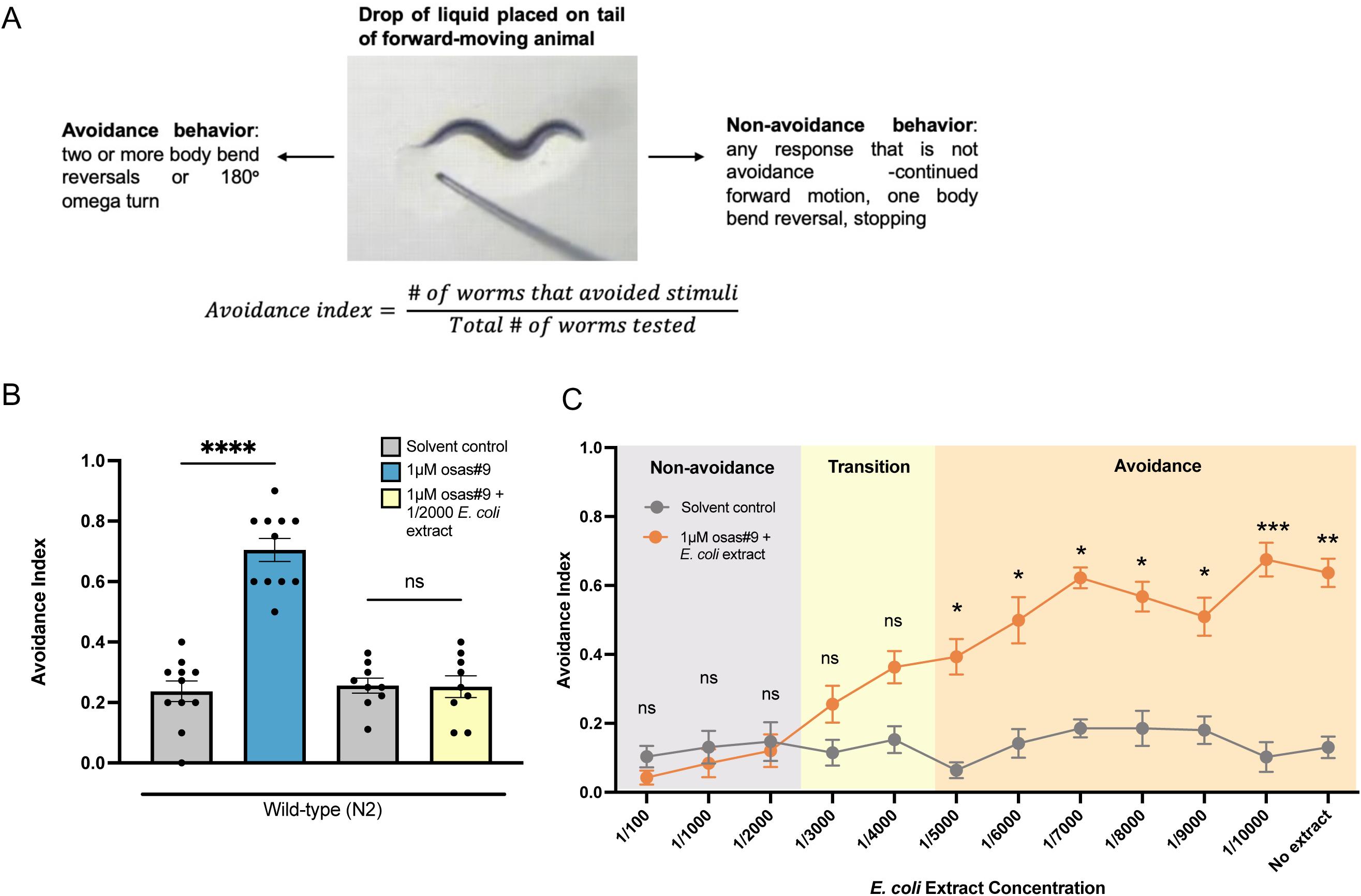
Food odor attenuates osas#9 avoidance in food-deprived animals. a) Schematic of avoidance index assay. Created with BioRender.com b) Food-deprived worms will no longer avoid 1µM osas#9 when *E. coli* extract is present in the stimulus mixture. n >= 9. c) Food extract induced osas#9 avoidance attenuation is dependent on extract concentration. When high concentrations (1/100-1/2000) of extract are present in the osas#9 mixture, avoidance is strongly attenuated. Intermediate concentrations of food extract (1/3000- 1/4000) still suppress avoidance but there is a trend of increasing avoidance. Low concentrations of food extract (1/5000-1/10000) present in the mixture are not sufficient to attenuate osas#9 avoidance. n >= 8. Error bars are SEM. *p < 0.05, **p < 0.01, ***p < 0.001, ****p < 0.0001, Shapiro-wilk normalcy test followed by ordinary one-way ANOVA w/ Sidak’s multiple comparisons (if data are normally distributed) or Kruskal-Wallis test w/ Dunn’s multiple comparisons (if data are not normally distributed).

### Food-dependent osas#9 avoidance attenuation is dependent on serotonin signaling

We wanted to know which neurotransmitters, if any, are involved in modulating the nervous system’s suppression of osas#9 avoidance in the presence of food extract. We tested *tph-1, cat-2*, and *tdc-1* mutants. *tph-1* encodes tryptophan hydroxylase, an enzyme required for the neuronal biosynthesis of serotonin (Sze et al., 2000; Yu et al., 2023). *cat-2* encodes an enzyme required for dopamine synthesis, tyrosine hydroxylase (Lints and Emmons, 1999). Finally, *tdc-1* mutants lack both tyramine and octopamine (Alkema et al., 2005). We found that *tph-1* mutants, while they avoided osas#9 alone normally (Figure S2A), continued to avoid osas#9 even when it was mixed with food extract (Figure 2A). This result indicates that serotonin is required for the integration of food odor cues into the osas#9 avoidance neural circuit. We also tested mutants that are deficient in glutamatergic synaptic transmission (*eat-4*) (Lee et al., 1999) and GABA biosynthesis (*unc-25*)(Jin et al., 1999), but these mutants did not avoid osas#9 alone normally, so those strains were excluded from our screen (Figure S2).

**Figure 2:**
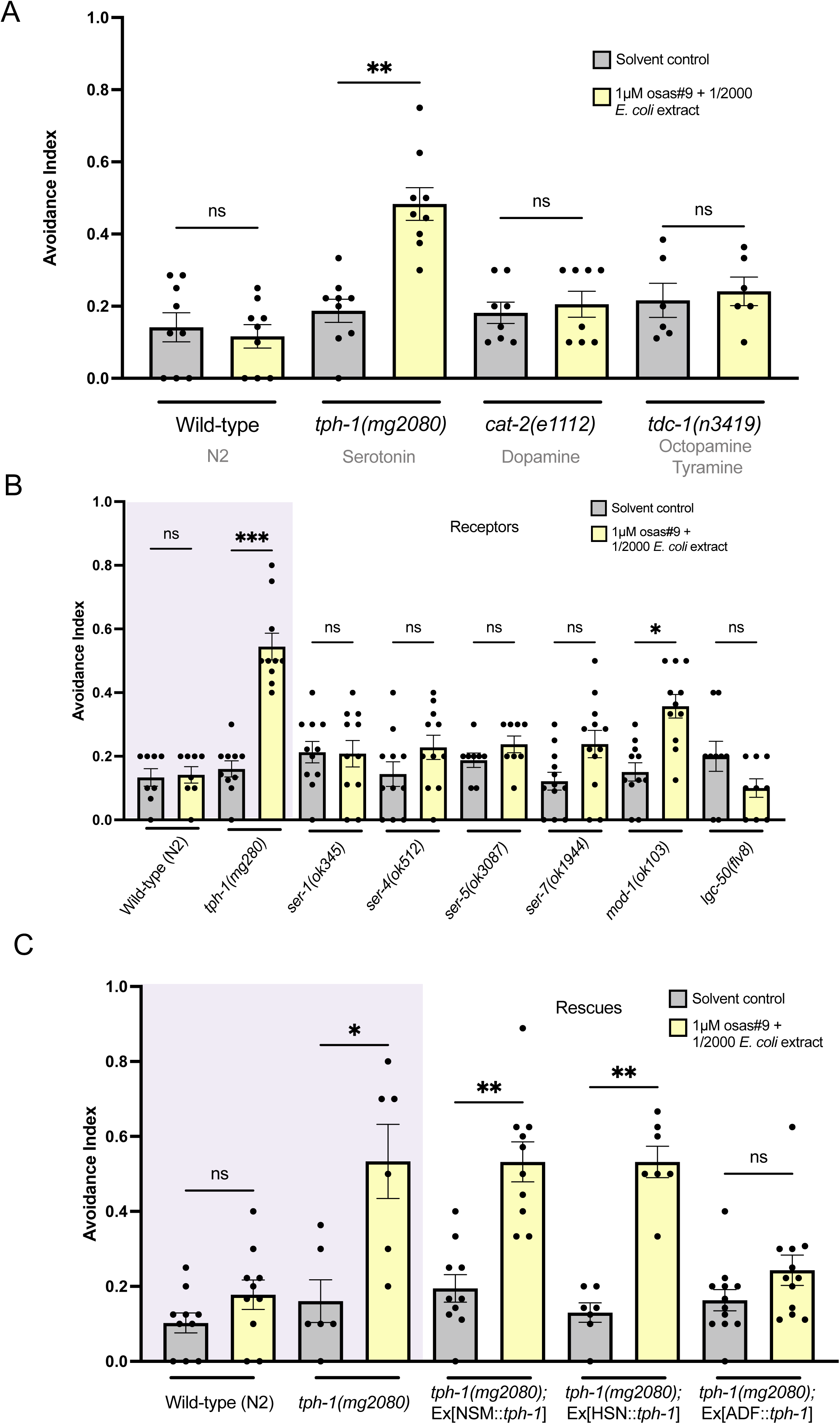
Serotonergic signaling is required for food-dependent osas#9 avoidance attenuation. a. A screen of neurotransmitter mutants showed that *tph-1(mg2080)* mutants avoid the osas#9 and food extract mixture. n >= 6. b. *mod-1(ok103)* mutants avoid osas#9 when mixed with food extract. n>=8. c. Rescuing *tph-1* expression in the ADF neurons restored wild-type osas#9 + food extract response. Rescuing *tph-1* in the NSM and HSN neurons resulted in no behavioral rescue. n>=6. Error bars are SEM. *p < 0.05, **p < 0.01, ***p < 0.001, ****p < 0.0001, Shapiro-wilk normalcy test followed by Kruskal-Wallis test w/ Dunn’s multiple comparisons.

There are six known serotonin receptors in *C. elegans* (Chase, 2007; Hapiak et al., 2009; Morud et al., 2021). To find out which serotonin receptor(s) were acting to attenuate osas#9 avoidance in the presence of *E. coli* extract, we tested mutants for all six serotonin receptors: *ser- 1, ser-4, ser-5, ser-7, mod-1*, and *lgc-50*. The *ser* receptors are reported to be GPCRs (Carre-Pierrat et al., 2006; Chase, 2007; Hapiak et al., 2009) whereas MOD-1 is a chloride channel (Ranganathan et al., 2000) and LGC-50 is a cation channel (Morud et al., 2021). We found that *mod-1* mutants avoided osas#9 mixed with food, indicating that this receptor is essential for the proper integration of the food cue (Figure 2B). However, we also noticed that avoidance behavior in *mod-1* worms trended a little bit lower that avoidance behavior of *tph-1* mutants.

In addition to investigating which serotonin receptors were acting to suppress osas#9 avoidance, we wanted to know which serotonin-releasing neuron(s) may be responsible for the avoidance suppression. There are three main neurons in *C. elegans* that express *tph-1*, and therefore, these are the three main neurons that produce serotonin: ADF, HSN, and NSM (Sze et al., 2000; Chang et al., 2006; Kullyev et al., 2010). There is also mild *tph-1* expression reported in the AIM and RIH neurons (Sze et al., 2000). The HSNs are primarily involved in egg-laying behavior (Weinshenker et al., 1995; Shyn et al., 2003); while the NSM neurons are implicated in the bacterial-slowing response (Sawin et al., 2000) and to a lesser extent, phalangeal pumping (Riddle DL, 1997; Li et al., 2012) – both food-related behaviors. The ADF neurons are unique in that they are the only serotonergic sensory neurons (Sze et al., 2000; Chang et al., 2006; Kullyev et al., 2010). Furthermore, the ADF neurons serve more diverse functions than other serotonergic neurons; they are involved in chemotaxis (Bargmann and Horvitz, 1991a), dauer entry (Bargmann and Horvitz, 1991b; Hu, 2005-2018), and sex-specific pheromone response (Fagan et al., 2018). We tested *tph-1* cell-specific rescues in ADF, HSN, and NSM neurons in a *tph-1* loss of function background. We found that rescuing *tph-1* expression in the ADF neurons restored wildtype food-dependent osas#9 avoidance attenuation (Figure 2C). To confirm *tph-1* expression in the ADF neurons we imaged animals that contained both a *tph-1* reporter (*tph-1*::GFP) and mCherry expression in the ADF neurons (*srh-142*::mCherry); we saw that the ADF neurons contained both proteins, confirming that *tph-1* is expressed in the ADF neurons (Figure S2B).

Overall, these results show that serotonin signaling from the ADF neurons plays an essential role in canceling the osas#9 avoidance signal in the presence of food odor.

### Calcium transients increase in the ADF neurons upon *E. coli* extract exposure

To better understand how the ADF neurons might be responding to food-derived chemicals, we performed calcium imaging. Transgenic worms expressing GCaMP in the ADF neurons under the *srh-142* promotor were loaded into microfluidic device (Reilly et al., 2017) and exposed to bacterial extract. We observed an increase in calcium transients in the ADF neurons upon stimulation with bacterial extract (Figure 3A). At 1/1000 and 1/2000 concentrations of extract, we saw a strong increase in calcium transients (Figure 3A-C; Figure S3A-B); this is consistent with the strong avoidance attenuation we observed at these concentrations of food extract (Figure 1C). We wanted to know if intermediate concentrations of bacterial extract created a response in the ADF neuron, so we stimulated the neuron with 1/3000 and 1/4000 concentration of food extract (Figure 3A,3D-E; Figure S3C-D). We saw that at 1/3000 concentration of food odor, the neuron showed a significant response (Figure 3D); while at 1/4000, the neuron did not significantly respond (Figure 3E). However, it should be noted that while the neuron did not significantly respond to food extract at the 1/4000 concentration, a heatmap of individual animal response shows a trend to weakly respond in 36% (4/11) of animals (Figure S3D) – although this observation is more subjective. This result showed us that these intermediate concentrations appear to be on the lower end of where worms will detectably still respond to food extract, consistent with behavioral data that shows weaker avoidance attenuation (Figure 1C). Finally, we hypothesized that low concentrations of bacterial extract, concentrations which were unable to attenuate osas#9 avoidance, would not produce any observable increase in calcium transients in the ADF neurons. To test this, we exposed animals to 1/8000 food odor (a low concentration chosen from behavioral data) and saw that there was no observable neural response (Figure 3A,3F; Figure S3E). These imaging experiments indicate that the ADF neurons have increased calcium transients in response to food odors when there is a sufficient amount of food, further supporting our behavioral data that osas#9 avoidance attenuation is dependent on food concentration.

**Figure 3:**
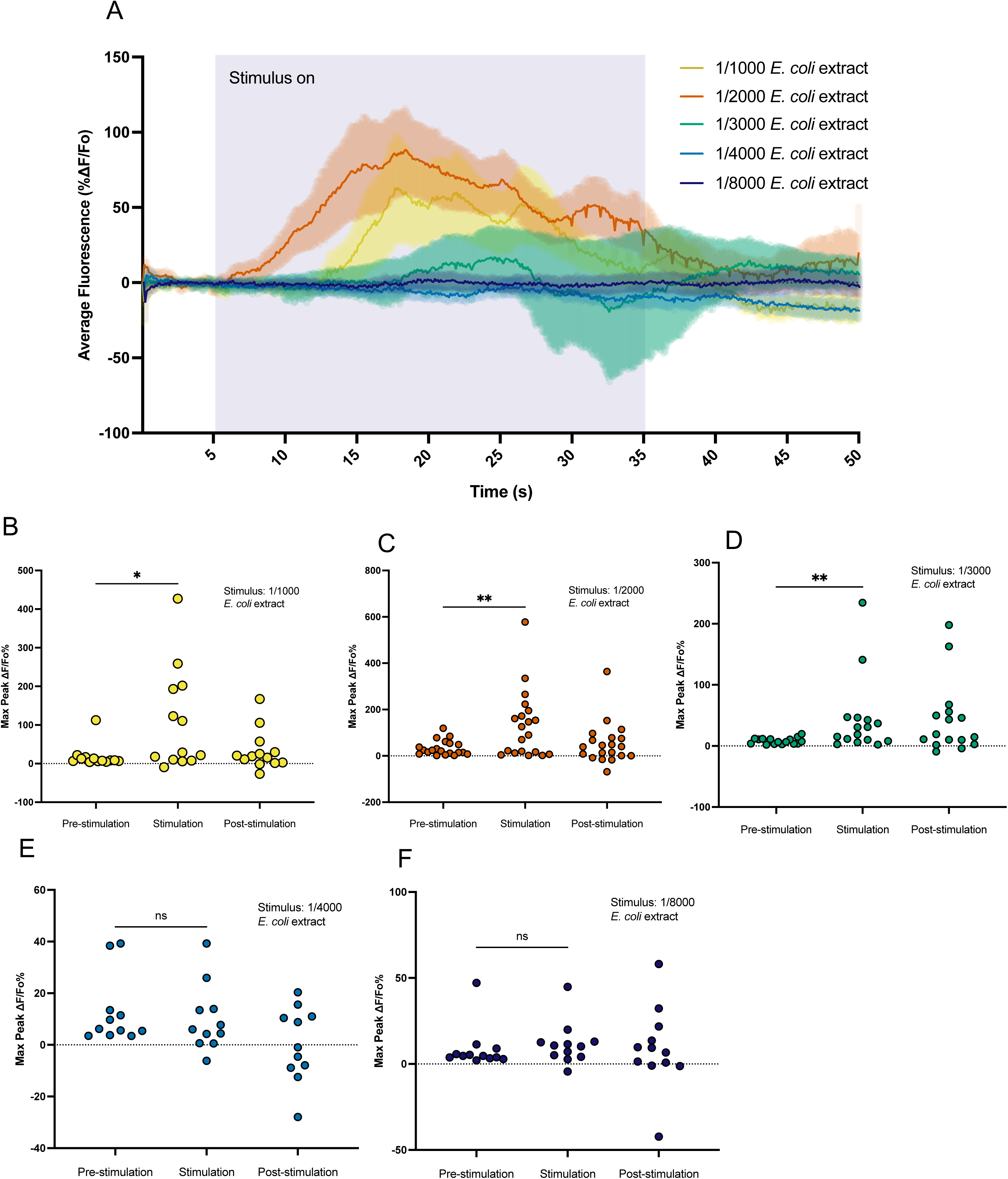
The ADF neuron responds to food odor. a. Calcium dynamics of the ADF neuron in ADF::GCaMP animals upon exposure to different concentrations of *E. coli* extract. Different concentrations depicted in different colors. Error clouds surrounding trace of means are SEM. n>=11. b. Maximum fluorescence intensity in the ADF neurons is significantly greater after exposure to 1/1000 food extract. n=13. c. Maximum fluorescence intensity in the ADF neurons is significantly greater after exposure to 1/2000 food extract. n = 20. d. Maximum fluorescence intensity in the ADF neurons is significantly greater after exposure to 1/3000 food extract. n>16. e. There is no difference in maximum fluorescence intensity in the ADF neurons pre and post exposure to 1/4000 food extract. n=11. f. There is no difference in maximum fluorescence intensity in the ADF neurons pre and post exposure to 1/10000 food extract. n=12. *p < 0.05, **p < 0.01, ***p < 0.001, ****p < 0.0001. Shapiro-wilk normalcy test followed by two-tailed Wilcoxon test pre stim vs. stim. Note that for b-f y-axis scales differ to best show data.

### The ADF neurons require input from other neurons in order to respond to food extract

Since we found that the ADF neurons are essential for integrating food odor signals into the nervous system, we wanted to find out if the ADF neurons require input from other cells to integrate the food signal. To answer this question, we crossed the ADF imaging strain into *unc- 13* and *unc-31* mutant backgrounds. These mutants are deficient in synaptic and peptidergic signaling, respectively (Richmond et al., 1999; Speese et al., 2007). If the ADF neurons still respond to food in the absence of synaptic and peptidergic communication, this will lead us to believe that the ADF neurons directly sense *E. coli* extract. However, if the ADF neurons requires synaptic or peptidergic signaling to respond, we will hypothesize that the ADF neurons function downstream of a different sensory neuron. Upon exposing either *unc-13* or *unc-31* ADF imaging strains to food extract, we saw that there was no significant increase in fluorescence in the ADF neuron upon food exposure in either mutant (Figure 4A-C; Figure S4A-B). However, examining individual animals revealed that 33% (4/12) of *unc-13* mutants responded, suggesting a trend of response for mutants that lack synaptic signaling (Figure S4A) and indicating that the ADF neurons may have some role in the absence of synaptic communication. These data suggest that the ADF neurons require both synaptic and peptidergic input from other neurons in order to fully respond to food extract.

**Figure 4:**
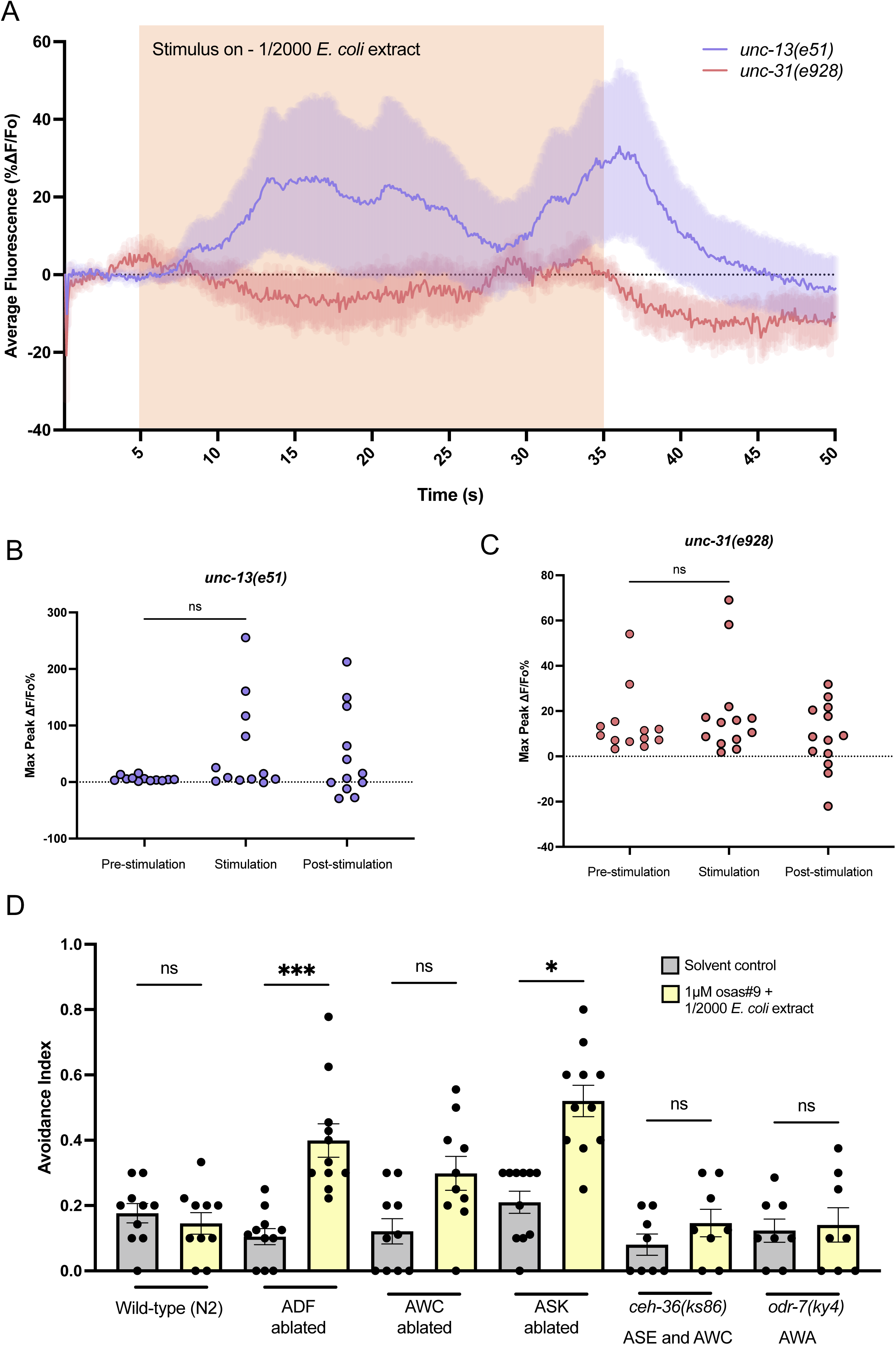
The ADF neurons require synaptic and peptidergic input for response to food extract. a. Calcium dynamics of the ADF neurons in ADF::GCaMP animals in *unc-13(e51)* and *unc-31(e928)* backgrounds upon exposure to 1/2000 *E. coli* extract. Error clouds surrounding trace of means are SEM. n=12. b. There is no difference in maximum fluorescence intensity in the ADF neuron pre and post exposure to 1/2000 food extract in an *unc-13(e51)* background. n = 12. Shapiro- wilk normalcy test followed by two-tailed Wilcoxon test pre stim vs. stim. c. There is no difference in maximum fluorescence intensity in the ADF neuron pre and post exposure to 1/2000 food extract in an *unc-31(e928)* background. n =12. Shapiro- wilk normalcy test followed by two-tailed Wilcoxon test pre stim vs. stim. d. Ablation of the ADF and ASK neurons results in avoidance to osas#9 + food extract. Ablation of the AWC neurons has no effect on avoidance attenuation behavior. *ceh- 36(ks86)* and *odr-7(ky4)* show normal avoidance attenuation. n >=8. Shapiro-wilk normalcy test followed by Kruskal-Wallis test w/ Dunn’s multiple comparisons. *p < 0.05, **p < 0.01, ***p < 0.001, ****p < 0.0001

We hypothesized that other neurons that are known to be involved in food sensation may be acting in the osas#9 avoidance attenuation neural circuit in concert with the ADF neurons.

Specifically, we theorized that a food-sensing neuron may be communicating with the ADF neurons. We tested ADF, ASK, and AWC neural ablation strains, in addition to *ceh-36* (defective sensation in ASE and AWC neurons (Koga and Ohshima, 2004)), and *odr-7* (defective sensation in AWA neurons (Sengupta et al., 1994)) mutants for their ability to suppress osas#9 avoidance in the presence of bacterial extract. Consistent with a requirement for *tph-1* expression in the ADF neurons, we observed that worms avoided osas#9 in the presence of bacterial extract when the ADF neurons were ablated (Figure 4D). In addition, we found that the ASK ablation strain avoided the osas#9-food mixture (Figure 4D). These results suggest that the ASK neurons may also act integrate food signals into the nervous system.

## Discussion

Like all animals, *C. elegans* are able to process multiple pieces of sensory information (Stein et al., 2014; Yau et al., 2015; Ghosh et al., 2017; Harris et al., 2019). During behavior, when a new cue is introduced, the neural signal from the new cue is incorporated into the nervous system, which can result in different behavioral output. We found that while food-deprived worms typically avoid the dispersal pheromone osas#9, they will no longer avoid this pheromone when it is mixed with an extract of the food (Figure 1b). Osas#9 avoidance attenuation is dependent on the strength of food extract present (Figure 1C), indicating that the nervous system weighs the strength of each cue when processing multiple pieces of information, this principle is true across the animal kingdom (Stein and Stanford, 2008; Leathers and Olson, 2012)

Through our work, we found that food-dependent osas#9 avoidance attenuation requires serotonergic signaling from the ADF neurons (Figure 2C). In humans, serotonin is associated with reward and punishment systems and thus plays a role in reward-related decision-making (Seymour et al., 2012). Depleted serotonin and mutations in serotonin receptors are associated with riskier decision-making (Rogers, 2011). Serotonin’s role in decision-making likely works through its ability to modulate neural circuits (Celada et al., 2013; Jacob and Nienborg, 2018; Marquez and Chacron, 2020). In multiple mammal models and across multiple sensory systems, serotonin application has been shown to dampen sensory neural signals (Jacob and Nienborg, 2018; Marquez and Chacron, 2020). And because serotonin receptors can be inhibitory or excitatory, there is a variety of ways that serotonin signaling can tune a circuit (Celada et al., 2013). Our results suggest that serotonin acts in a similar neuromodulatory role in *C. elegans* when animals encounter food extract during sensation of the aversive pheromone osas#9.

We found that the inhibitory serotonin receptor MOD-1 was required for food-dependent osas#9 avoidance attenuation (Figure 2B). MOD-1 is expressed in numerous interneurons and motor neurons that modulate worm movement (Harris et al., 2009; Hammarlund et al., 2018).

Since MOD-1 is an inhibitory chloride channel (Ranganathan et al., 2000), we propose that during food sensation, MOD-1 is activated on avoidance-promoting neurons, repressing avoidance behavior. Since we also noticed that *mod-1* mutants seemed to avoid the osas#9 and extract mixture less strongly than the *tph-1* mutants, there is a possibility some other serotonin receptors may play smaller roles in integrating food cues into the nervous system. However, their role(s) may be too small to detect when MOD-1 is functioning, as it appears to be the main receptor. Alternatively, there is some research that suggests there could be as many as eight serotonin receptors in *C. elegans,* leaving the possibility that an uncharacterized receptor acts in the food odor integration pathway (Carre-Pierrat et al., 2006).

Moreover, we show that the ADF neurons require synaptic and peptidergic input from other neuron(s) for their response to food extract (Figure 4A-C). From a screen of food-related neurons, we believe that the ASK neurons are sending input to the ADF neurons (Figure 4D). However, there may be other neurons involved as our screen of neurons was not exhaustive.

There are numerous receptors, including neuropeptide receptors, expressed on the ADF neurons that could be screened in the future to narrow down the nature of this signaling (Hammarlund et al., 2018). Given that the ADF neurons require other neural input to respond to food extract, we suggest that although the ADF is a sensory neuron, it is functioning as an interneuron in this instance. This is consistent with some prior research that has shown that although the ADF neurons are sensory, they can sometimes function as interneurons (Guo et al., 2015).

Overall, we propose a model in which, upon sufficient *E. coli* extract sensation, an inhibitory neural pathway is activated to suppress an avoidance neural pathway. We suggest that the ASK neuron senses food extract and excites the ADF neurons. The ADF neurons send serotonergic signals that are received via MOD-1 receptor (Figure 5). The MOD-1 receptor is expressed on movement modulating interneurons such as AIZ, AIB, AIY, and AIA (Harris et al., 2009; Hammarlund et al., 2018). Prior research has shown that ablation of the AIZ and AIB interneurons causes a reduction in reversals, implying that these interneurons promote reversal and avoidance behavior (Tsalik and Hobert, 2003; Bono and Villu Maricq, 2005). Given that the inhibitory MOD-1 receptor is expressed on both of these neurons, it is possible that in the presence of food extract, serotonergic signaling from the ADF neurons inhibits reversal interneurons that are usually activated in the presence of osas#9. This type of coordinated inhibition is seen across neural circuits in the animal kingdom (Yan, 2002; Wang, 2008; Miller et al., 2011; Wu and Guo, 2011; Chadderton et al., 2014; Jacob and Nienborg, 2018; Marquez and Chacron, 2020; Herd et al., 2021). Our work adds to the body of evidence that serotonin serves an important function during multi-sensory processing.

**Figure 5:**
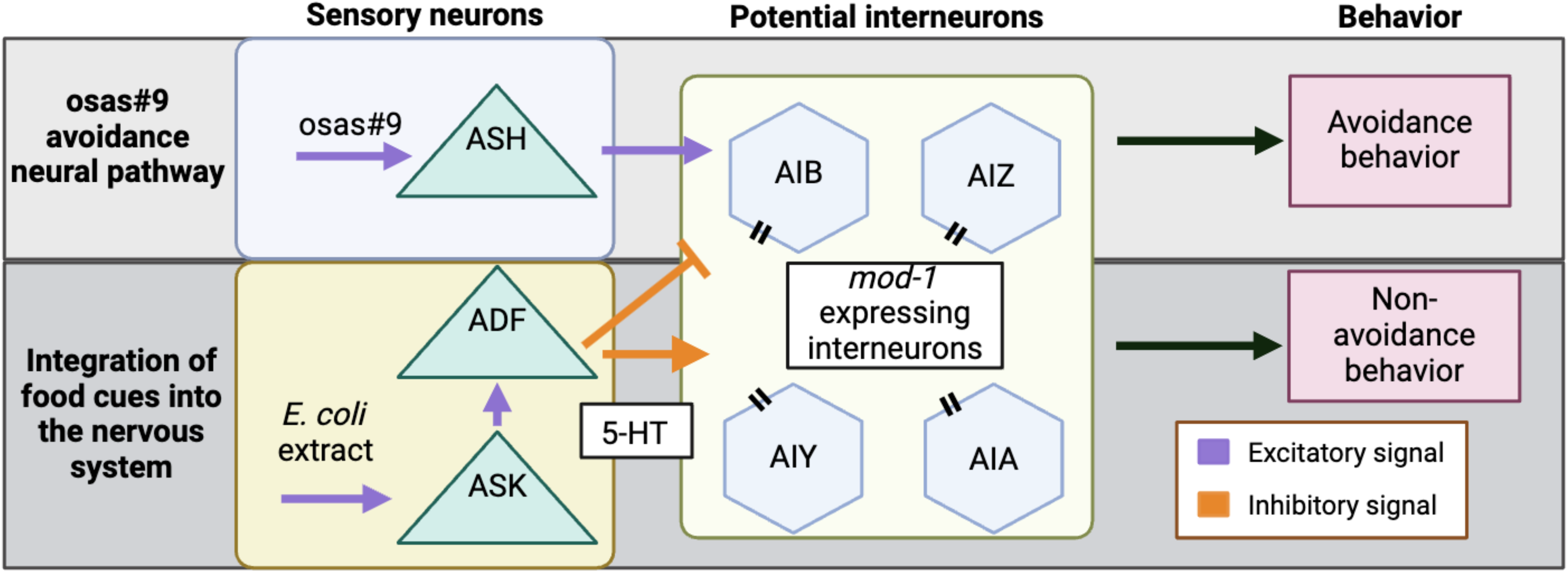
Proposed model for osas#9 avoidance attenuation. osas#9 is sensed primarily by the ASH sensory neurons (Chute et al., 2019). The ASH neurons communicate with downstream interneurons neurons to initiate avoidance. When animals are exposed to food extract, the ADF neurons send serotonergic signals that inhibit the osas#9 avoidance pathway. The serotonergic signal sent by the ADF is likely received via MOD-1 receptors. MOD-1 receptors are expressed on interneurons that are known to modulate forward and backward motion. Created with BioRender.com

## Acknowledgements

We thank the Portman, Bargmann, Flavell, Sternberg, and Ferkey labs for generously sharing worm strains. Some strains were also provided by the CGC, which is funded by NIH Office of Research Infrastructure Programs (P40 OD010440). We also thank all current and past members of the Srinivasan lab for their continuous feedback on this project. Specifically, we thank Chris Chute for pioneering parts of this project. Funding: R01DC016058

## Supplementary figure legends

**Table S1:**

| Strain name | Genotype | Source |
| --- | --- | --- |
| N2 | Wildtype | CGC |
| MT15434 | *tph-1 (mg280)* II | CGC |
| CB1112 | *cat-2(e1112)* II | CGC |
| MT13113 | *tdc-1(n3419)* II | CGC |
| CX13503 | *eat-4 (ky5)* | Bargmann Lab |
| CB156 | *unc-25(e156)* III | CGC |
| DA1814 | *ser-1(ok345)* X | CGC |
| AQ866 | *ser-4(ok512)* III | CGC |
| RB2277 | *ser-5(ok3087)* I | CGC |
| RB1585 | *ser-7(ok1944)* X | CGC |
| MT9668 | *mod-1(ok103)* V | CGC |
| SWF912 | *lgc-50(flv8)* III; *flvIs2[tph-1p(short)::Chrimson + elt-2p::mCherry]* | Flavell Lab/CGC |
| CX13571 | *tph-1(mg280)*; *kySi56*; *kyEx4077= srh-142::nCre* (95 ng/uL); *myo-3::mCherry* (5ng/uL) | Bargmann Lab |
| CX13572 | *tph-1(mg280)*; *kySi56*; *kyEx4057= ceh-2::nCre* (10 ng/uL); *myo-3::mCherry* (5ng/uL) | Bargmann Lab |
| SWF855 | *tph-1(mg280)*; *flvIs2[tph-1(NSM-specific fragment)::Chrimson, elt-2::mCherry]*; *flvEx401[pegl-6::tph-1 cDNA, pmyo-2::mcherry]* | Flavell lab |
| JSR187 | *mgIs42 [tph-1::GFP + rol-6(su1006)]*; *vlcIs10 [srh-142::mCherry + elt-2::GFP]* | Srinivasan lab |
| NFB1445 | *rrf-3(pk1426)* II; *lite-1(ce314)* X; *vlcEx1292[srh-142p::mCherry::SL2::GCaMP3 + unc-122p::RFP]* | CGC |
| JSR182 | *unc-31 (e928)*; *rrf-3(pk1426)* II; *lite-1(ce314)* X; *vlcEx1292 [srh-142p::mCherry::SL2::GCaMP3 + unc-122p::RFP]* | Srinivasan lab |
| JSR184 | *unc-13 (e51)*; *rrf-3(pk1426)* II; *lite-1(ce314)* X; *vlcEx1292 [srh-142p::mCherry::SL2::GCaMP3 + unc-122p::RFP]* | Srinivasan lab |
| UR987 | *him-5(e1490)* V; *udEx212[Pelt-2:: GFP; Psrh-142::GFP; Psrh-142::ced-3(p15); Psrh-142::ced-3(p17) line 2]* | Strain from Portman lab and extrachromosomal array generated in Ferkey lab:<br>https://doi.org/10.1371/journal.pgen.1006153 |
| PS6022 | qrIs1[*sra-9::mCasp1*] | Sternberg lab |
| PY7502 | oyIs85 [*ceh-36p::TU#813* + *ceh-36p::TU#814* + *srtx-1p::GFP* + *unc-122p::DsRed*] | CGC |
| FK311 | *ceh-36(ks86)* X | CGC |
| CX4 | *odr-7(ky4)* X. | CGC |

**Figure S1:**
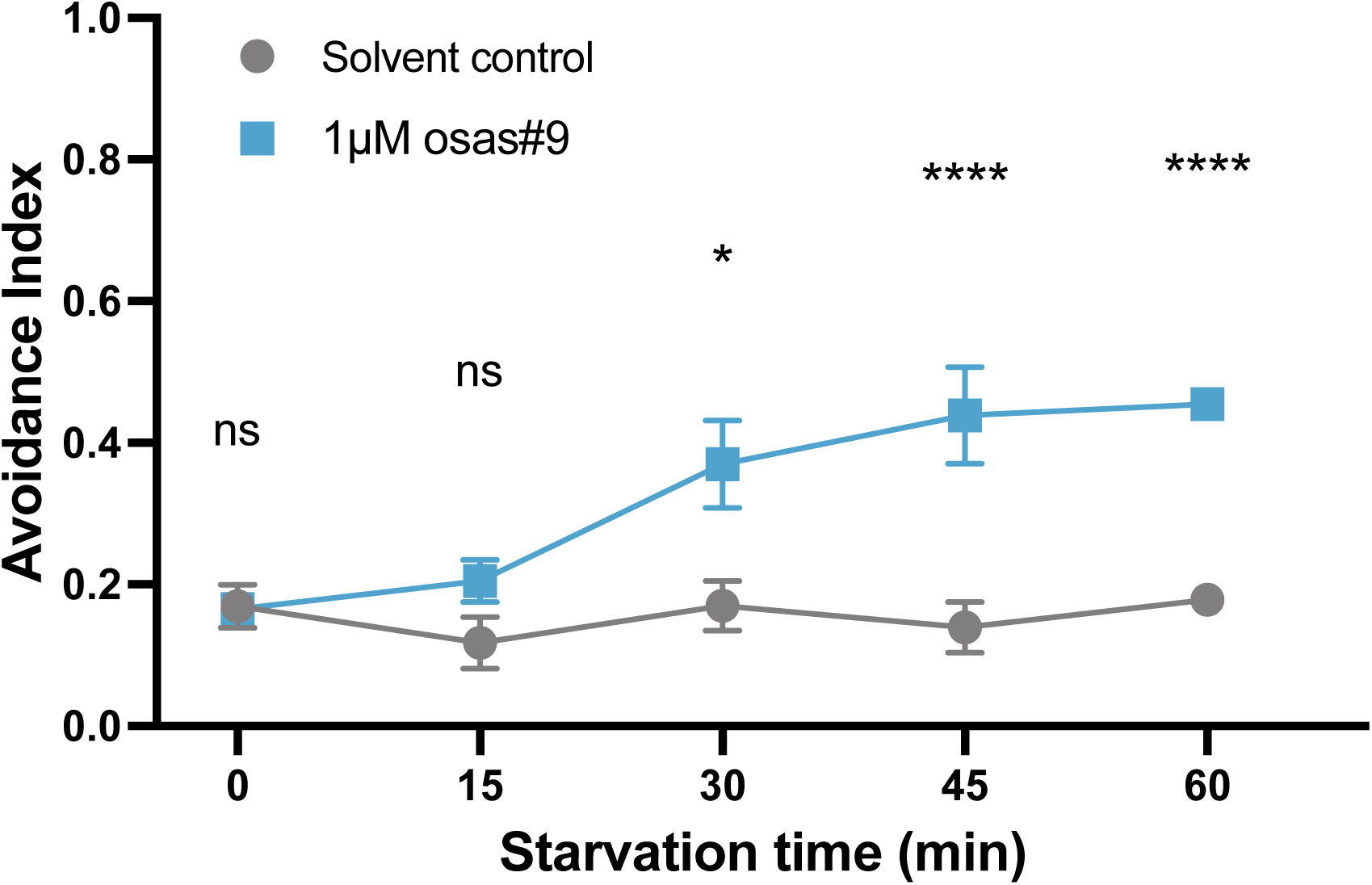
Animals will begin avoiding 1µM osas#9 30 minutes after food removal. Prior to 30 minutes, avoidance index to 1µM osas#9 is no different than the solvent control. The avoidance index appear to plateau after 30 minutes of food removal. n >= 7.

**Figure S2:**
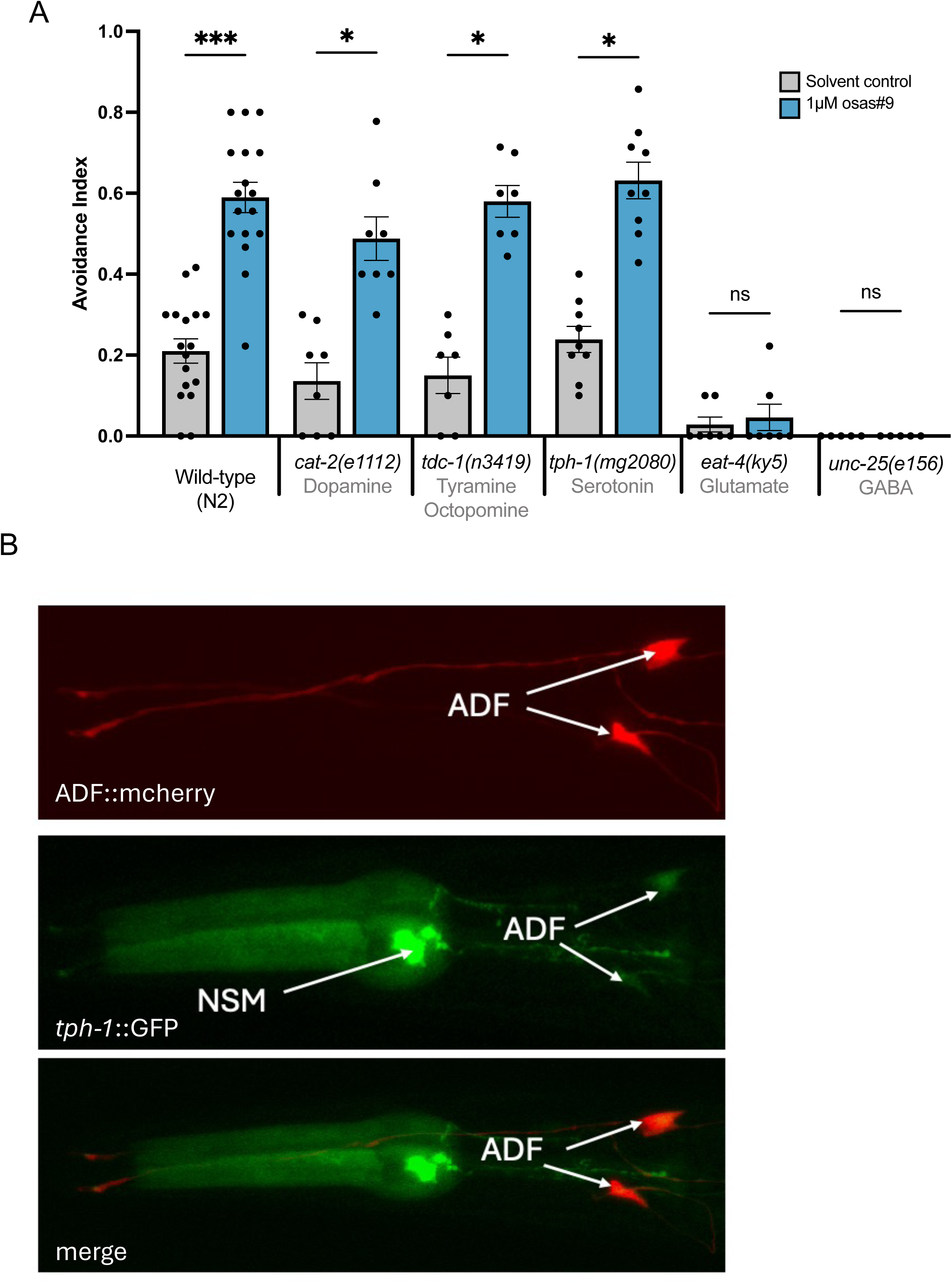
a. Glutamate and GABA are required for osas#9 avoidance A screen of neurotransmitter mutants showed that *eat-4(ky5)* mutants and *unc-25(e156)* do not avoid osas#9. n >= 5. Error bars are SEM. *p < 0.05, **p < 0.01, ***p < 0.001, ****p < 0.0001, Shapiro-wilk normalcy test followed by Kruskal-Wallis test w/ Dunn’s multiple comparisons. b. *tph-1* is expressed in the ADF neurons Animals containing both *tph-1*::GFP and ADF(*srh-142)*::mCherry constructs showed overlapping red and green expression in the ADF neurons. 63X magnification; oil objective

**Figure S3:**
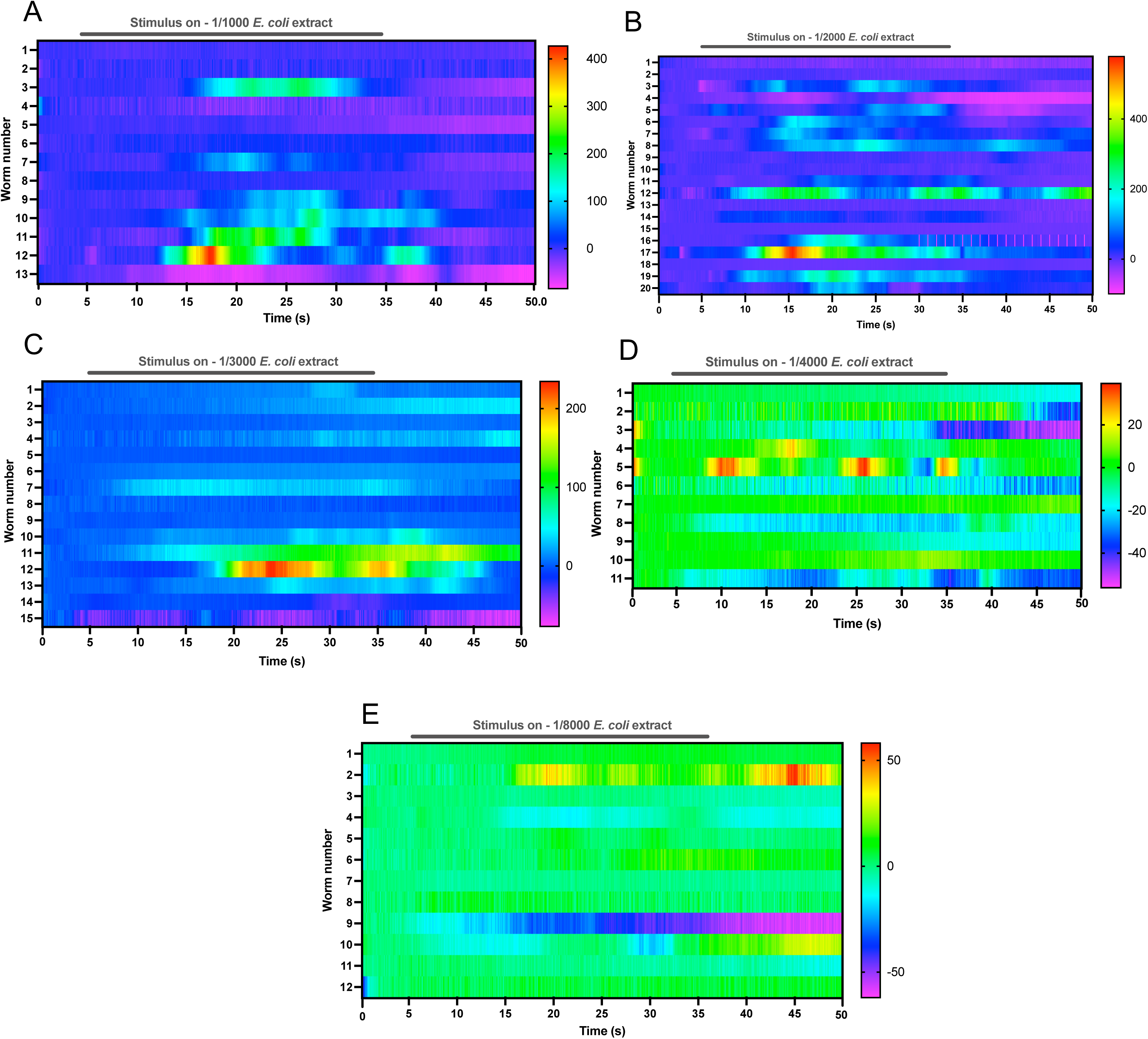
Heatmaps of fluorescence intensity in the ADF neurons in ADF::GCaMP animals upon exposure to a.1/1000 *E. coli* extract, b. 1/2000 *E. coli* extract, c. 1/3000 *E. coli* extract, d. 1/4000 *E. coli* extract, e. 1/8000 *E. coli* extract. n >=11.

**Figure S4:**
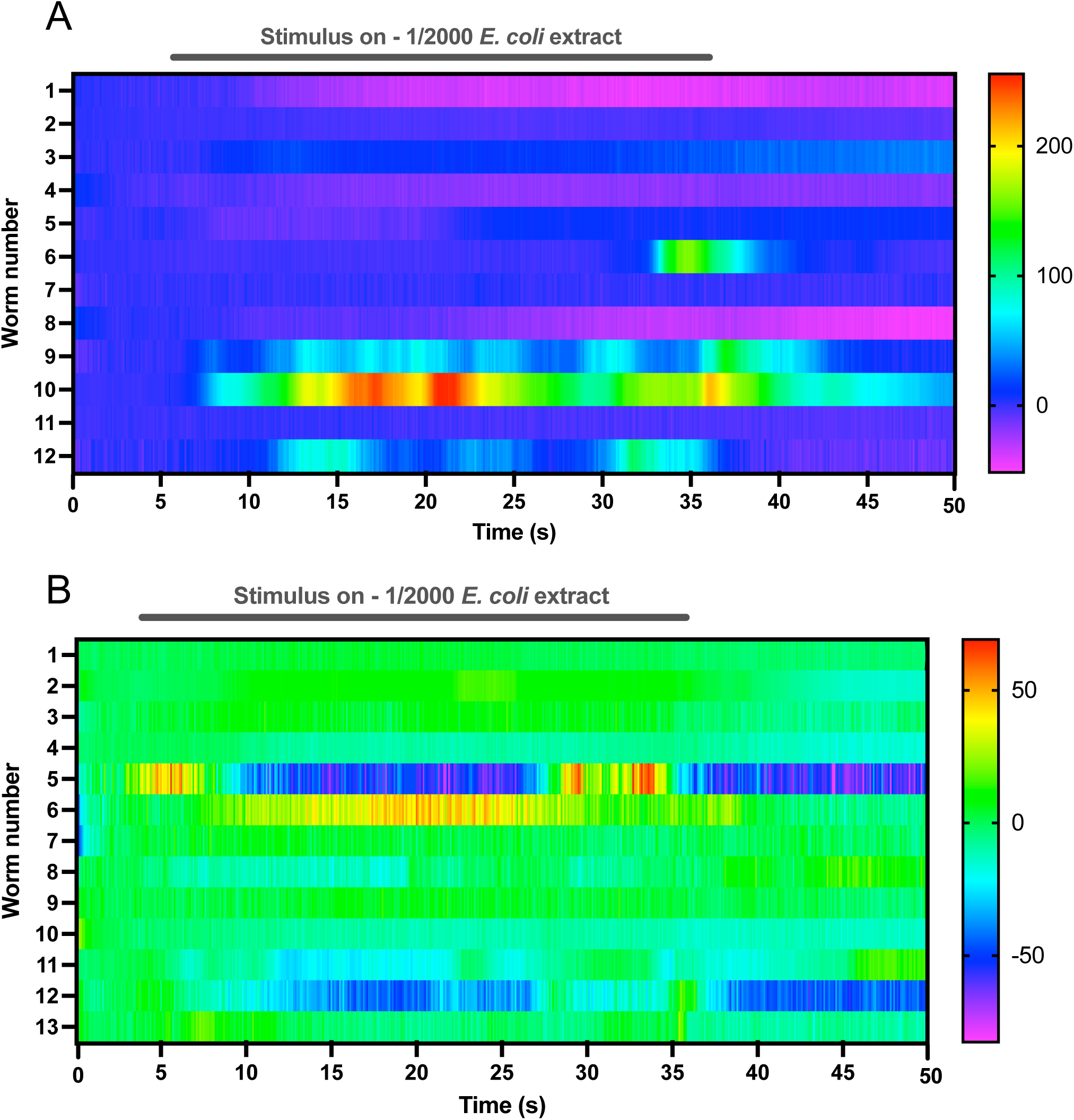
Heatmaps of fluorescence intensity in the ADF neuron in ADF::GCaMP animals upon exposure to 1/2000 *E. coli* extract in a. an *unc-13(e51)* background and b. an *unc-31(e928)* background.

